# B-type plexins regulate mitosis via RanGTPase

**DOI:** 10.1101/2024.01.21.576518

**Authors:** Nicholus Mukhwana, Ritu Garg, Magali Williamson

## Abstract

Aberrant mitosis can result in aneuploidy and cancer. The small GTPase, Ran, is a key regulator of mitosis. B-type Plexins regulate Ran activity by acting as RanGTPase activating proteins (GAPs) and have been implicated in cancer progression. However, whether B-type plexins have a role in mitosis has not so far been investigated.

We show here that PlexinB1 functions in the control of mitosis. Depletion of PlexinB1 resulted in defects in chromosomal segregation, increasing cells with multipolar spindles and prolonging metaphase, while activation of PlexinB1/B2 with Sema4D produced aberrant spindles.

An increase in MAD1-positive kinetochores in PlexinB1/B2 depleted cells indicated application of spindle assembly checkpoint (SAC), possibly resulting from faulty spindle-microtubule assembly, and PlexinB1 depletion promoted regrowth of acentrosomal microtubules and defects in spindle pole refocussing. Depleted cells that had bypassed SAC possessed lagging chromosomes and chromosomal bridges.

The mitotic defects observed upon PlexinB1 depletion were rescued by an RCC1 inhibitor, indicating that PlexinB1 signals, at least in part, via Ran to affect mitosis.

These errors in mitosis generated multinucleate cells, and nuclei of altered morphology and abnormal karyotype. Furthermore, Sema4D-treatment increased the percentage of cells with micronuclei, precursors of chromothripsis. Defects in B-type plexins may contribute to the well-established role of plexins in cancer progression by inducing chromosomal instability.

## Introduction

Frequent mis-segregation of chromosomes at anaphase during mitosis, known as chromosomal instability leads to aneuploidy and more aggressive forms of cancer^1^. Chromosomal misegregation can also lead to chromothripsis^2^, fragmentation of chromosomal regions resulting in multiple rearrangements, promoting cancer progression.

The small GTPase, Ras-related nuclear protein (Ran), is a key controller of mitosis ^3^. Ran cycles between a GTP-bound active form and a GDP-bound inactive form^4^. RanGDP is converted to RanGTP by its chromatin-bound guanine exchange factor (GEF), Regulator of chromosome condensation 1 (RCC1),^5^ while RanGAP1, catalyses the intrinsic GTPase activity of Ran converting RanGTP to RanGDP, ^6^ ^7^. The hydrolysis of RanGTP by RanGAP1 is stimulated by RanBP1 which binds to RanGTP^5^ ^8^and inhibits RCC1.

RanGTP is thought to function by inducing the localised release of regulatory factors from inhibitory complexes made between karyopherins (exportins and importins)^9^ and cargo proteins during nuclear trafficking in interphase cells or spindle assembly factors during mitosis, which contain nuclear localisation sequences (NLS) or nuclear export sequences (NES)^4^.

During mitosis, chromatin-bound RCC1 generates a gradient of RanGTP around the chromosomes ^10^ and RanGTP in the vicinity of the chromosomes induces the localised release of importin-bound spindle assembly factors (SAFs) from importins.^11,12^ Ran has a role in microtubule stabilisation and nucleation close to the chromosomes during mitosis and in acentrosomal spindle formation. ^13,14^

Some RanGTP also occurs in the centrosomes and regulates centrosomal cohesion, microtubule-nucleation and duplication licensing.^15^

Due to its key role in mitosis and nucleocytoplasmic translocation, Ran and its regulators have been implicated in many cancers. ^16^ For example, loss of RanGAP1 expression promotes osteosarcoma.^17^

B-type plexins are also regulators of Ran, promoting the conversion of RanGTP to RanGDP by acting as RanGTPase activating proteins (GAPs). ^18^ Depletion of either PlexinB1 or PlexinB2 increase the levels of RanGTP in the cell, while activation of PlexinB1 or B2 by their ligand Sema4D, decreases RanGTP levels^18^. Vertebrates possess nine plexin genes (class A(1-4), B(1-3) C1 and D1),^19^ acting as receptors for twenty or more semaphorins. Plexins and semaphorins have been implicated in many cancers, both as tumour suppressors and metastasis-promoting genes.^20^

PlexinB1, the most well studied of the class B plexins, is the receptor for semaphorin 4D (Sema4D) ^21^ and Sema3C.^22^ PlexinB1 regulates nucleocytoplasmic trafficking of androgen receptor through direct activation of the GTPase activity of Ran. However, whether B-type plexins have a role in mitosis has not so far been investigated/reported.

We questioned whether PlexinB1 has a role in the regulation of mitosis through its activity as a RanGAP.

## Results

### 1 PlexinB1 has a role in the regulation of mitosis

RanGTP has a role in the regulation of mitotic spindle formation.^3^ RanGTP also controls the localised release of spindle assembly factors from inhibitory complexes made with importins^11,12^ and correct balance of RanGTP/GDP is required for proper spindle assembly.^3^ High concentrations of RanGTP around the chromosomes, resulting from RCC1 bound to the chromatin, induces non-centromeric microtubule nucleation and centrosome-free spindle MT growth.^23^ ^24^ To determine whether PlexinB1 affects these Ran-associated phenotypes, PlexinB1 levels were knocked down with siRNA and the effect on mitosis examined.

Hela cells expressing GFP-tubulin and mCherry-Histone were transfected with non-silencing control siRNA or siRNA to PlexinB1 and the number of cells with multipolar spindles were analysed. Spindle poles were identified with pericentrin staining. PlexinB1 depletion resulted in a significant increase in the number of dividing Hela cells with multipolar spindles and with three or more centrosomes (Figure 1A). An increase the number of dividing cells with multipolar spindles and aberrant spindle formation was also found upon depletion of PlexinB1 in NP1542 cells, cells derived from non-transformed prostate tissue^25^ (Figure 1B).

**Figure 1.**
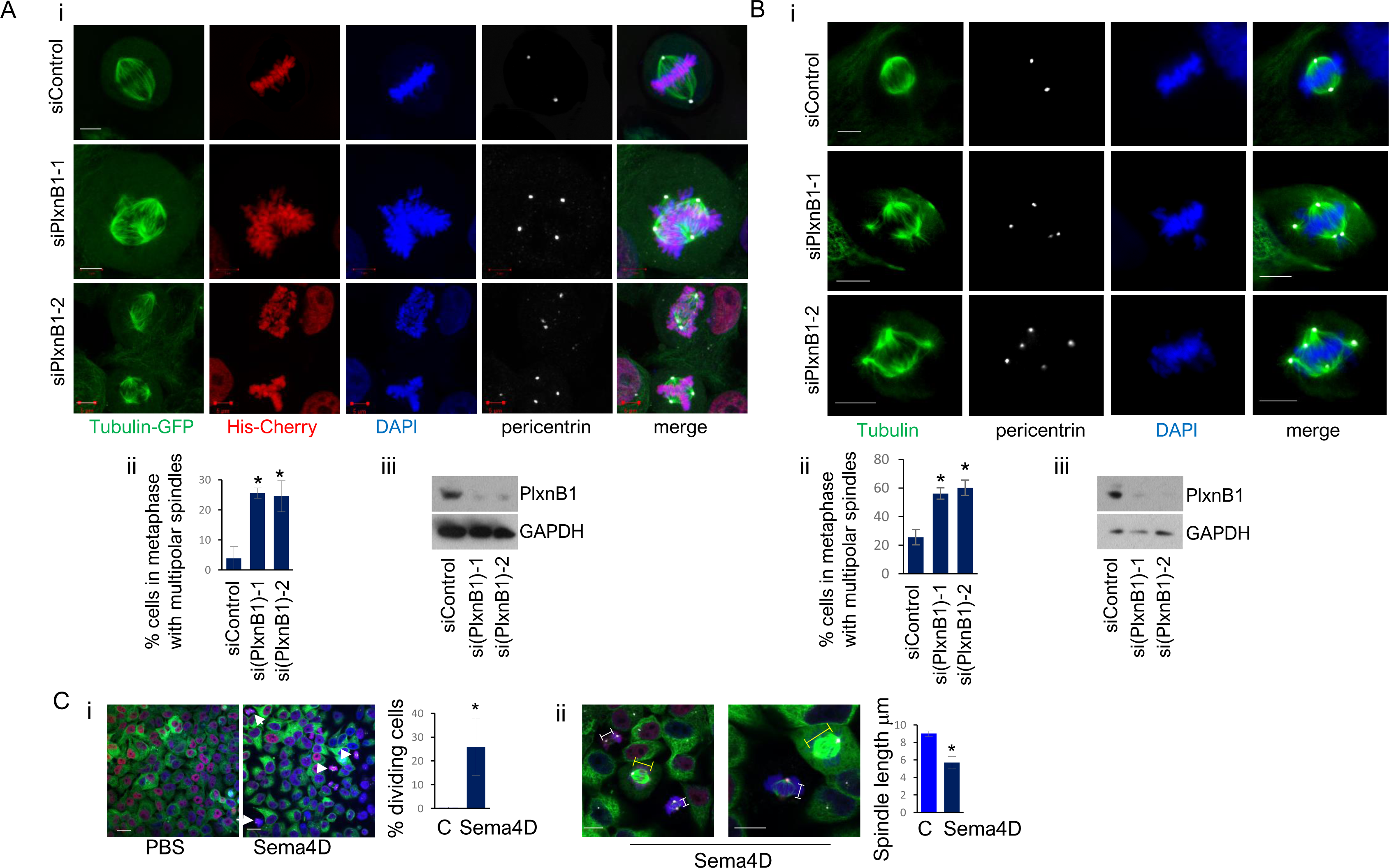
Depletion or activation of PlexinB1 affects mitotic spindle formation. **A.** i). Representative images of HeLa cells expressing GFP-tubulin and mCherry-Histone transfected with control siRNA or two different siRNAs against PlexinB1, stained with DAPI and pericentrin. Scale bar, 5 μm. ii). Bar chart depicts the % of dividing cells with multipolar spindles marked by pericentrin staining (n=3, minimum of 91 dividing cells scored per condition*p=<0.05, 2 tailed student T-Test). iii). Western blot analysis of knockdown of PlexinB1 by siRNA in Hela cells. **B.** i). Representative images of NP1542 cells transfected with control siRNA or two different siRNAs against PlexinB1, stained with DAPI and pericentrin. Scale bar, 5 μm. ii). Bar chart depicts the % of dividing cells with multipolar spindles marked by pericentrin staining (20+ cells in metaphase scored per condition per experiment, n=3, *p=<0.05, 2 tailed student T-Test, vs control iii). Western blot analysis of knockdown of PlexinB1 by siRNA in NP1542 cells. **C i).** Representative images of HeLa cells expressing GFP-tubulin and mCherry-histone, treated with Sema4D-Fc (2μg/ml) or vehicle control for 6 h, fixed and stained for DAPI and pericentrin (magnification x40, scale bars=20 μm). Arrows show cells with collapsed spindles; bar chart shows % of cells with collapsed spindle (see also Supplementary Figure 1). **ii)**. HeLa cells expressing GFP-tubulin and mCherry-histone treated with Sema4D-Fc (2μg/ml) for 6 h, stained for DAPI and pericentrin (magnification x63, scale bars=10 μm) White and yellow bars indicate length of spindle for dividing cells. Bar chart shows average spindle length of cells in metaphase treated with Sema4D or vehicle for 6 h (average of dividing 10 cells measured).

Furthermore, activation of PlexinB1 and/or PlexinB2 by treatment of cells with Sema4D resulted in an increase in dividing cells with collapsed spindles with a shortened spindle axis (Figure 1C, Supplementary Figure 1).

Defects in mitosis frequently results in the operation of a spindle assembly checkpoint (SAC) which delays mitosis until all kinetochores are attached to the mitotic spindle.^26^ Aberrant mitosis consequently results in a pausing of the cycle at metaphase until SAC is overcome. In order to examine whether plexinB1 depletion has any influence on the timing of mitosis, time lapse videos were recorded of Hela cells in which PlexinB1 expression had been knocked down by siRNA. Depletion of PlexinB1 in HeLa cells significantly increased the length of time spent to proceed from prometaphase to anaphase as measured by time lapse video microscopy (Figure 2A, supplementary videos). Depletion of PlexinB1 significantly increased the percentage of dividing cells that took over 170 minutes to reach anaphase from prometaphase (Figure 2B). Consistent with these results, PlexinB1 depletion also increased the percentage of cells in metaphase at fixed time points (Figure 2C). Depleted cells displaying a defect in exiting metaphase were frequently subjected to apoptosis eventually, observed by time lapse video microscopy and shown by activated caspase 3/7 staining (Figure 2D). This may be a response to mitotic catastrophe resulting from prolonged mitotic arrest caused by the activation of SAC.

**Figure 2.**
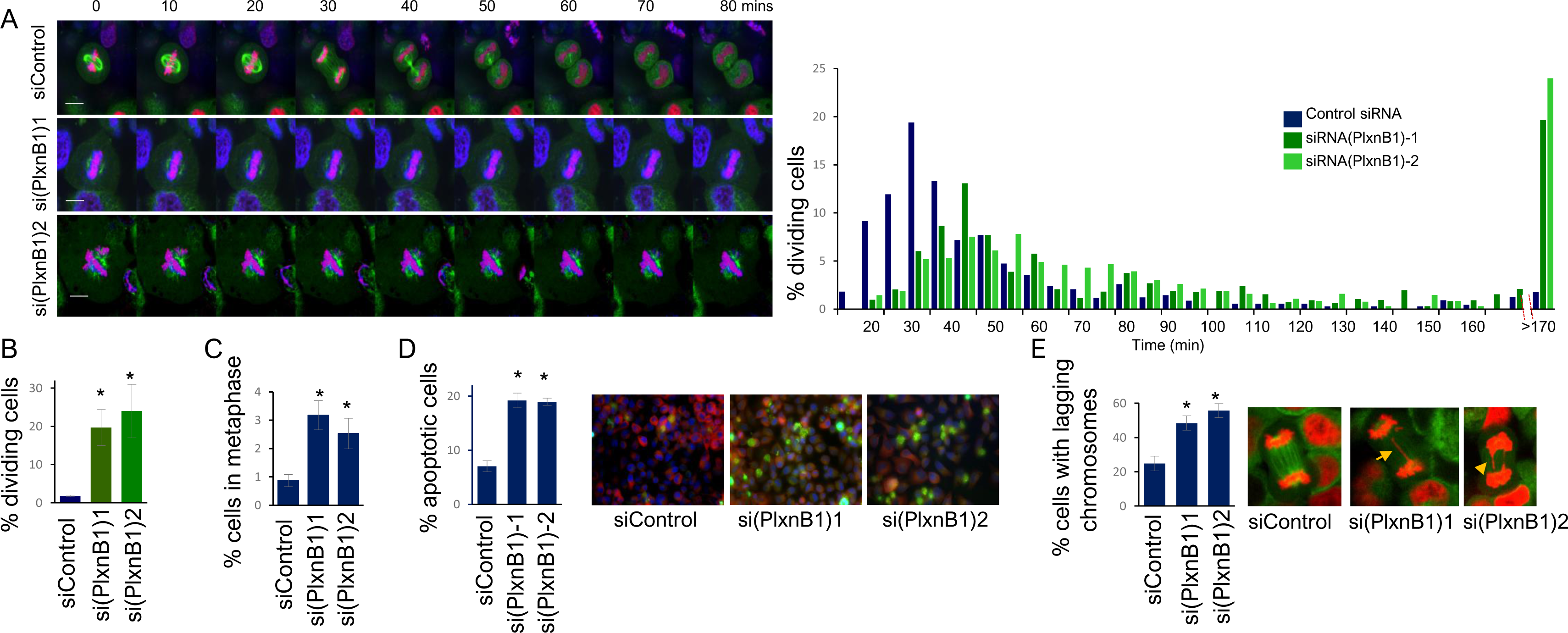
Effect of PlexinB1 depletion on mitosis. A.Depletion of PlexinB1 delays mitosis. Representative images from time lapse video of HeLa cells expressing GFP-tubulin and mCherry-Histone, transfected with non-silencing siRNA (siControl) or two different siRNAs against PlexinB1, cultured with NucBlue Live ReadyProbes Reagent, blue. (x60 magnification, scale bar= 10μm). Images were taken every 10 min, (n=3). Bar chart depicts the length of time for cells in prometaphase to reach anaphase. HeLa cells expressing GFP-tubulin and mCherry-Histone were transfected with non-silencing siRNA or two different siRNAs against PlexinB1 and the time taken for prometaphase to anaphase was scored from time lapse videos, images taken every 5mins for 15hrs . Data presented as the % of cells in mitosis that took the indicated time to reach anaphase from prometaphase. (>170 indicates % of dividing cells that took longer than 170 minutes for prometaphase to anaphase). Average of 3 experiments with 100+ cells scored for each experiment per condition. **B.** % of cells in mitosis where the time taken for prometaphase to anaphase exceeded 170min. (Average of 3 experiments with 100+ cells scored for each experiment per condition *p=<0.05, 2 tailed student T-Test) **C. Depletion of PlexinB1 expression increases the number of cells in metaphase**. HeLa cells expressing GFP-tubulin and mCherry-Histone plated on coverslips were transfected with non-silencing siRNA or two different siRNAs against PlexinB1. The cells were fixed, stained for DAPI and the number of cells in metaphase scored. (n=4, minimum of 365 cells scored per experiment per condition, *p=<0.05, 2 tailed student T-Test). **D. Depletion of PlexinB1 promotes apoptosis**. HeLa cells expressing GFP- tubulin, transfected with control or two different siRNAs against PlexinB1 were fixed and stained for activated Caspase-3/7 (representative images, magnification x40, scale bar = 20μm). Bar chart shows average number of cells staining positive for activation of caspase-3/7A, (n=3, *p=<0.01, 2 tailed student T-Test). **E. Depletion of PlexinB1 increases the number of cells in anaphase with lagging chromosomes**. HeLa cells expressing GFP-tubulin and mCherry-Histone transfected with non-silencing siRNA or two different siRNAs against PlexinB1. The % of dividing cells in anaphase with lagging chromosomes was scored from time lapse videos. (*p=<0.05, 2 tailed student T-Test). Average of 3 experiments with 50+ cells scored for each experiment per condition. Representative images of cells in anaphase, arrow depicts chromosomal bridge, arrowhead depicts lagging chromosome

To further investigate the effect of PlexinB1 depletion on the duration of mitosis, Hela cells, in which PlexinB1, B2 or RanBP1 expression had been knocked down, were synchronised by treatment with 10 µM RO-3306 or vehicle control for 20 hours, which arrests cells at the G2/M phase border. ^27^ The percentage of cells in the various stages of mitosis at fixed time points following the release from the G2/M block was recorded. Consistent with the time lapse results, depletion of either PlexinB1 or PlexinB2 prolonged the time in prometaphase (supplementary Figure 2). At 30mins there was a greater percentage of cells in prometaphase and a lower percentage of cells in metaphase in depleted cells vs control. After 60min, the majority of cells were in metaphase in control cells, while the majority were still in prometaphase in dividing cells in which PlexinB1 or PlexinB2 had been knocked down by siRNA. Knockdown of RanBP1, an activator of RanGAP1, used as a control, prolonged mitosis to a similar extent (supplementary Figure 2).

Furthermore, for cells that reached anaphase, and had bypassed SAC, loss of PlexinB1 expression significantly increased the percentage of cells in anaphase with lagging chromosomes or chromosomal bridges (Figure 2E). The increase in lagging chromosomes may be a consequence of merotelic attachment to the spindle ^28^

Together these results show that PlexinB1 has a role in the regulation of correctly orchestrated mitosis.

### 2. PlexinB1 depletion increases MAD1 staining at kinetochores

We have found that depletion of PlexinB1 prolongs the time taken for Hela cells to reach anaphase. Prolonged mitosis can result from the application of a spindle assembly checkpoint (SAC), ^29^ which postpones anaphase until all kinetochores are engaged.

To investigate whether PlexinB1/B2 depletion influences kinetochore-spindle attachment, synchronised cells were stained for MAD1, which binds to unattached kinetochores only. Depletion of either PlexinB1 or B2 resulted in a significant increase in MAD1 staining in synchronised cells, 30min after release from a G2/M block with RO-3306 (Figure 3A). Prolonged prometaphase delay was also found for RanBP1 depletion, used as a control. ^30^

**Figure 3.**
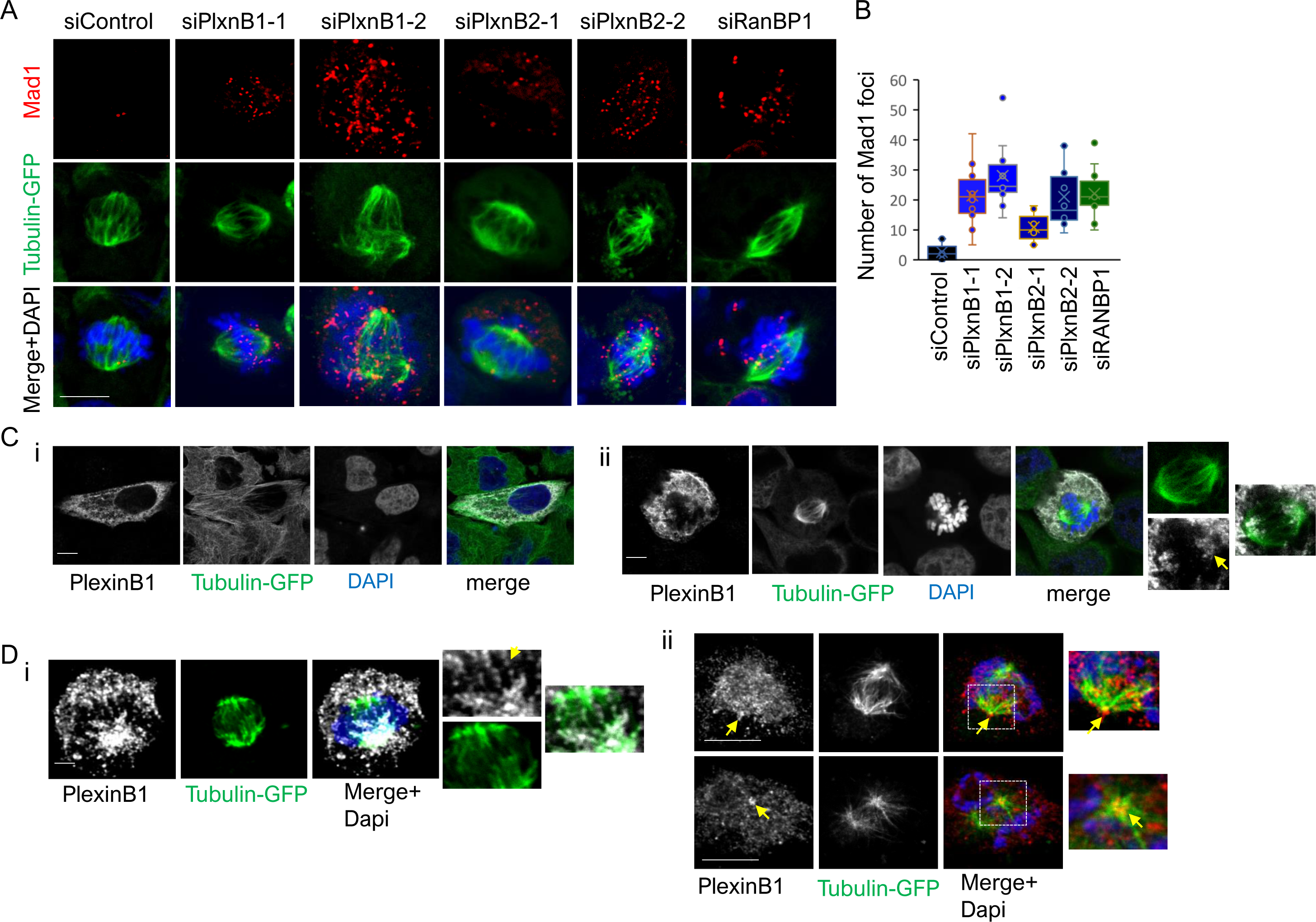
B-type Plexin depletion increases unattached kinetochores. A). HeLa cells expressing GFP-tubulin and mCherry-Histone were transfected with non-silencing siRNA or two different siRNAs against PlexinB1, PlexinB2 or against RanBP1. Cells were stained for Mad1 and the number of Mad1 spots counted, 7 or more dividing cells counted per condition. Scale bar = 10μm **B).** Box and whisker plot of the number of Mad1 foci for each condition. **C). Subcellular localisation of PlexinB1** Hela cells expressing tubulin-GFP transfected with PlexinB1(full-length)-myc, stained for Myc (PlexinB1), representative confocal image of interphase cell (i) and dividing cell (ii), arrow depicts PlexinB1 within the spindle structure in zoomed image. Single slices of z stack scale bar = 10μm **D).** HeLa cells expressing GFP-tubulin and mCherry-histone transfected with PlexinB1(FL)-myc, permeabilised with digitonin, stained for myc and Dapi (x63 magnification, scale bar=10 μm) i). arrow shows PlexinB1 expression close to central region of spindle. ii). arrow shows PlexinB1 expression around spindle poles.

These results imply that the delay in mitosis seen upon PlexinB1 depletion results from application of SAC. SAC overactivation is also seen for RanGAP1 knockdown ^17^(gong)

### 3 Localisation of PlexinB1 during mitosis

Plexins have a transmembrane domain and function as cell surface receptors. However, we have found that PlexinB1 affects nucleocytoplasmic trafficking by acting as a RanGAP, suggesting it has an intracellular function also. We next investigated where PlexinB1 is expressed in interphase and dividing cells. In interphase cells transfected with PlexinB1(FL)-myc, PlexinB1 is expressed in the cytoplasm as well as the plasma membrane (Figure 3C i,^18^). In dividing cells, PlexinB1 is localised to an area within the body of the mitotic spindle (Figure 3Cii, supplementary Figure 3A-C). In dividing cells permeabilised with digitonin, PlexinB1 expression localises to the spindles or kinetochores showing a similar distribution to that of RanGAP1^31,32^ within the spindle (Figure 3D i). In contrast to RanGAP1 ^31^ however, PlexinB1 also localizes at the immediate vicinity of spindle poles (Figure 3D ii).

### 4. PlexinB1 depletion affects spindle microtubule regrowth following microtubule disassembl

A delay in kinetochore attachment and subsequent application of SAC may be a consequence of aberrant spindle microtubule stabilisation or microtubule nucleation. High levels of RanGTP occur around chromosomes due to the association of active RCC1 with chromatin. A high level of RanGTP around chromosomes can induce microtubule nucleation and spindle formation in the absence of centrosomes. ^33^ Chromatin-mediated microtubule nucleation also contributes to early spindle assembly in the presence of centrosomes. ^24^

To assess the effect of PlexinB1 depletion on anastral microtubule nucleation, microtubules were disrupted by cold treatment for 30 mins during which time the MTs are almost completely disrupted (Figure 4A) and microtubule initiation and reassembly assessed at various time points after transfer to 37°C. After 3min reassembly, the majority of non-silenced cells had developed bipolar asters organised by centrosomes, with little sign of anastral microtubule polymerisation. Depletion of PlexinB1 resulted in a significant increase in the percentage of mitotic cells with acentrosomal microtubules after 3min (Figure 4B) and a decrease in mitotic cells with bipolar centrosomal asters only. By 20min a bipolar spindle was restablished for the majority of both silenced and non-silenced mitotic cells. An increase in the percentage of cells with non-focussed spindle poles was observed for PlexinB1 depleted cells after 20min incubation at 37°C (Figure 4C).

**Figure 4.**
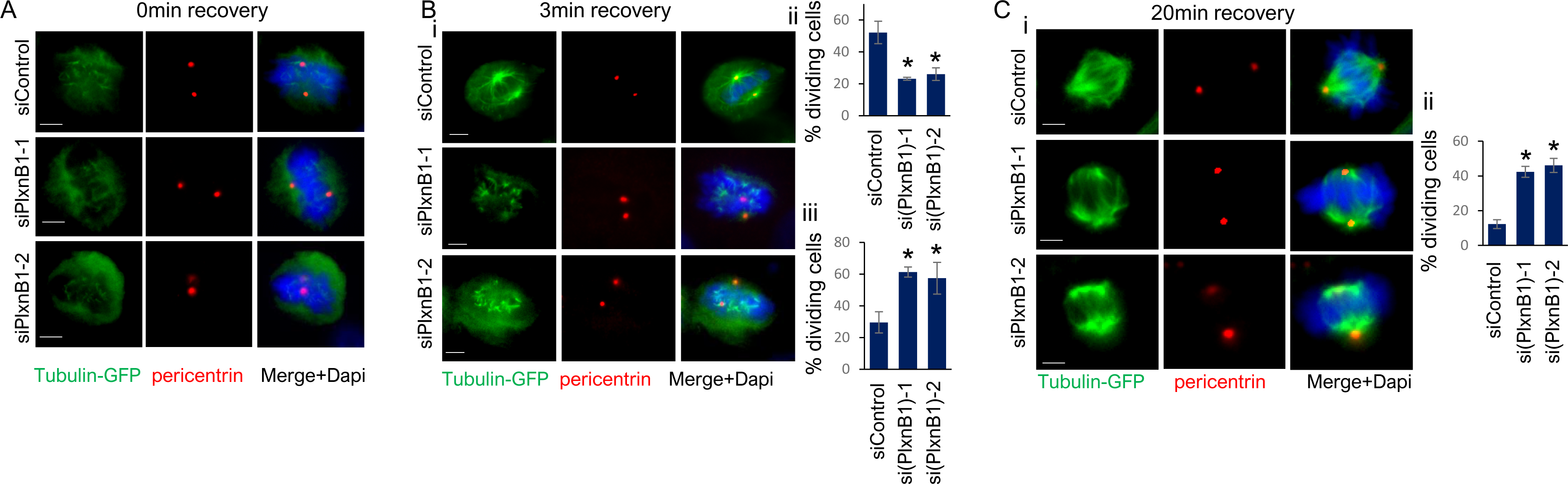
Effect of PlexinB1 depletion on spindle microtubule regrowth. Hela cells expressing tubulin-GFP and transfected with non-silencing siRNA (NS) or siRNA to PlexinB1, were kept on ice for 30mins. Cells were then placed in medium at 37°C for time indicated to allow microtubule regrowth, before fixation. Cells were stained for pericentrin. **A)** 0min recovery, representative images **B).** i). Representative images of microtubule regrowth assays after 3min recovery at 37°C ii) percentage of dividing cells with bipolar asters only (n=3, minimum of 20 metaphase cells scored per condition per experiment) error bars = SEM, (*p=<0.05, 2 tailed student T-Test) iii). percentage of dividing cells with acentrosomal asters. (n=3, minimum of 20 metaphase cells scored per condition per experiment) error bars = SEM, (*p=<0.05, 2 tailed student T-Test) **C)** i). Representative images of microtuble regrowth assays recovery after 20mins. ii). Graph shows percentage of dividing cells with non-focussed spindle poles (n=3, minimum of 16 metaphase cells scored per condition per experiment) error bars = SEM, (*p=<0.01, 2 tailed student T-Test) x40magnification, scale bar = 5μm

These results suggest that PlexinB1 has a role in control of non-centrosomal mitotic spindle assembly.

### 5. Mitotic defects resulting from PlexinB1 depletion are restored by inhibition of RCC1

We have found that PlexinB1 or PlexinB2 depletion affects the fidelity and timing of mitosis. PlexinB1 and B2 act as RanGAPS, catalysing the conversion of RanGTP to RanGDP. Ran has a key role in the regulation of mitosis and depletion of RanGAP1 results in aberrant mitosis.

We questioned whether PlexinB1/B2 exerted its effects on mitosis through Ran. Ran is activated by RCC1 (the GEF for Ran) which is active when chromatin bound. Methylation of RCC1 by protein arginine methyltransferase (PRMT) is required for the association of RCC1 with chromatin and the activation of Ran^34^. Treatment of cells with EPZ020411, a small molecule inhibitor of PRMT, results in a decrease in RanGTP levels^34^. To establish whether the contribution of PlexinB1 to the regulation of mitosis is dependent on Ran activity, PlexinB1 depleted cells were treated with RCC1 inhibitor and the effect on mitosis examined.

Treatment of PlexinB1-depleted HeLa cells with EPZ020411 significantly reduced the time taken for cells to reach anaphase from prometaphase, restoring the time taken to control levels (Figure 5A). Furthermore, treatment of PlexinB1-depleted cells with the RCC1 inhibitor significantly reduced the percentage of mitotic cells with multipolar spindles and with three or more centrosomes (Figure 5B).

**Figure 5.**
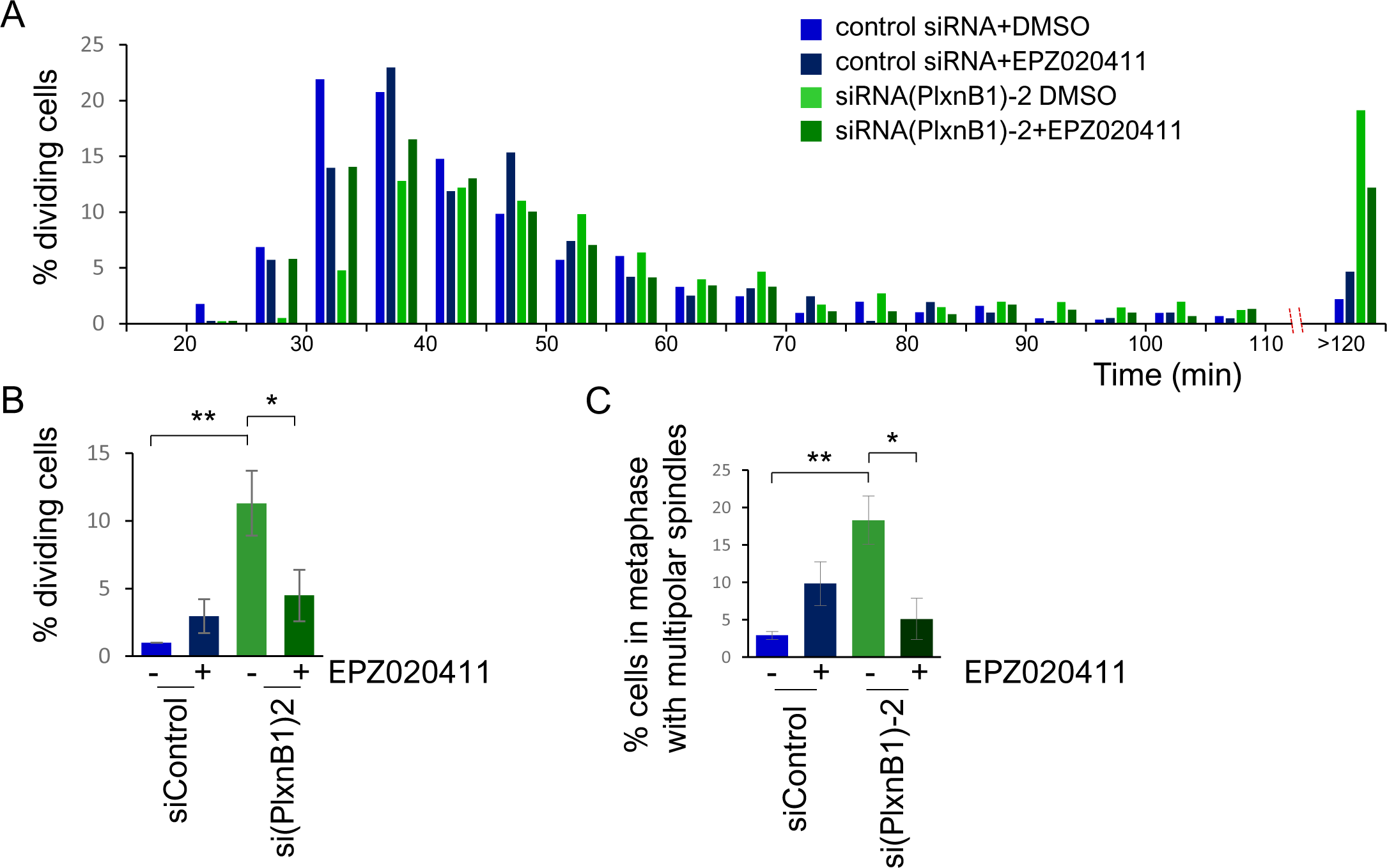
**RCC1 inhibition restores PlexinB1 depletion-induced mitotic defects. A). Inhibitor of RCC1 rescues time taken for prophase to anaphase**. HeLa cells expressing GFP-tubulin and mCherry-Histone were transfected with non-silencing siRNA or two different siRNAs against PlexinB1 and treated with EPZ020411 (50μM for 12 hr) or vehicle. The length of time for cells in prometaphase to reach anaphase was scored from time lapse videos, images taken every 5mins for 15hrs. Average of 4 experiments with 93+ cells scored for each experiment per condition. **B)**.Graph of the % of cells in mitosis where the time taken for prometaphase to anaphase exceeded 110min. (**p=<0.01 (control siRNA+DMSO vs siRNA(PlxnB1)+DMSO), * p=<0.05 siRNA(PlxnB1)+DMSO vs siRNA(PlxnB1)+EPZ020411, 2 tailed or 1 tailed student T-Test respectively). Average of 4 experiments with 93+ cells scored for each experiment per condition. Error bars denote SEM. **C)**. **Inhibitor of RCC1 rescues number of dividing cells with multipolar spindles.** % cells in metaphase with multipolar spindles. Average of 3 experiments with 100+ cells scored for each experiment per condition (**p=<0.01 (control siRNA+DMSO vs siRNA(PlxnB1)+DMSO), * p=<0.05 siRNA(PlxnB1)+DMSO vs siRNA(PlxnB1)+EPZ020411, 2 tailed student T-Test).

Together these results show that PlexinB1 plays a part in the regulation of correctly coordinated mitosis through Ran, consistent with its role as a RanGAP.

### 6. PlexinB1 depletion and activation drives chromosome mis-segregation

Defective mitosis can lead to cells with nuclear abnormalities, chromosomal instability and aneuploidy, common hallmarks of cancer. Multinucleation often results from defects in spindle formation and aberrant chromosome segregation.

To determine whether B-Type plexins influence chromosome stability, we examined the number and morphology of nuclei in cells transfected with control or PlexinB1 or PlexinB2- specific siRNA. Knockdown of expression of both PlexinB1 and PlexinB2 in Hela cells resulted in a significant increase in multinucleate cells (Figure 6A). Furthermore, knockdown of PlexinB1 or PlexinB2 had a significant effect on nuclear morphology as assessed by nuclear FormFactor, a measure of nuclear shape (Figure 6B ).^35^ Knockdown of PlexinB1 also increased the number of cells with two or more nuclei in NP142 cells (Figure 6C).

**Figure 6.**
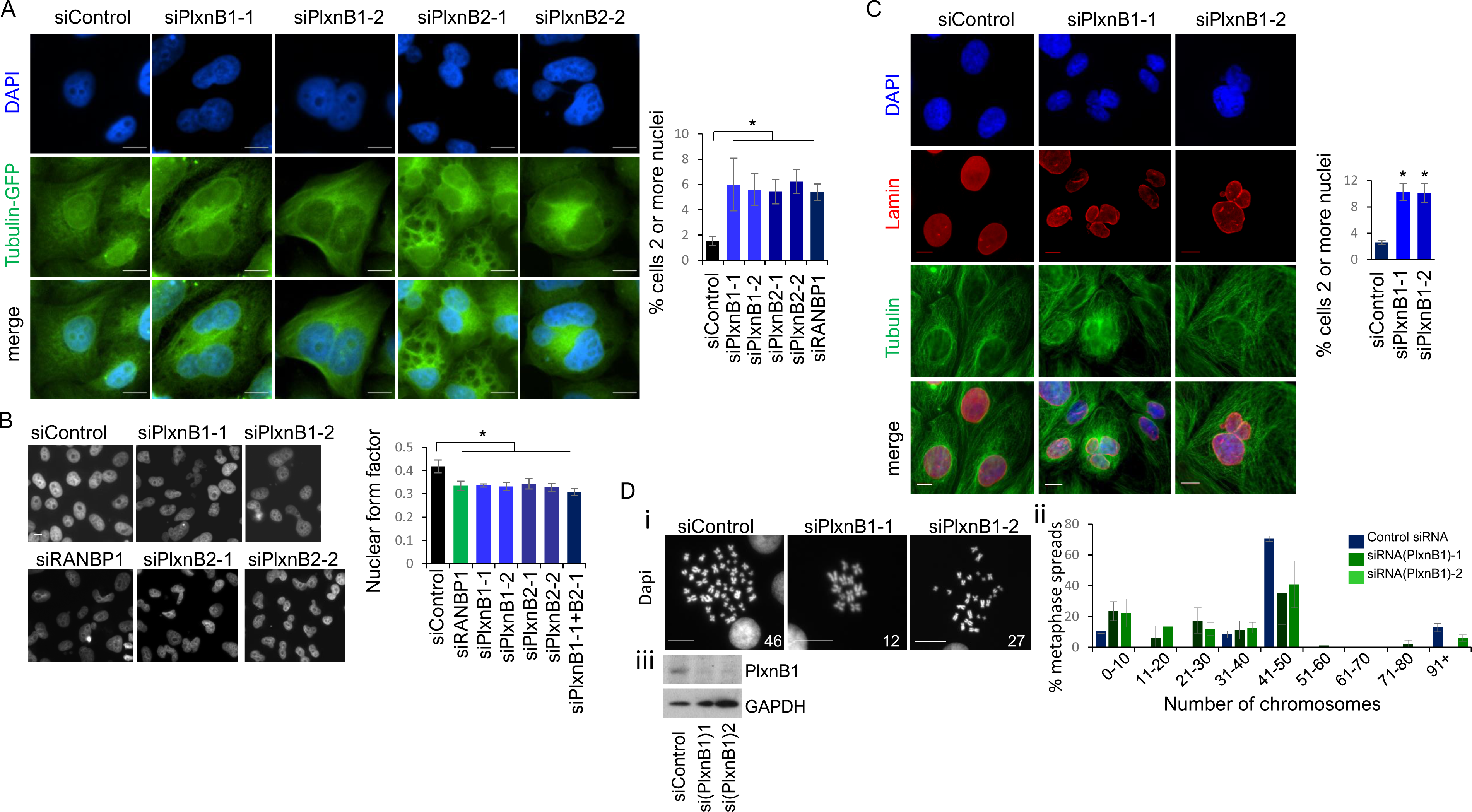
Effect of depletion of B-type plexins on chromosomal segregation. A). Representative images of HeLa cells expressing GFP-tubulin and mCherry-Histone, transfected with control siRNA or two different siRNAs against PlexinB1 or PlexinB2, stained with DAPI, scale bar =10μm. Graph shows % of Hela cells with two or more nuclei in cells transfected with control siRNA or siRNA to PlexinB1, PlexinB2 or RanBP1 (minimum of 3,000 cells scored per condition over 4 experiments, *p=<0.05, 2 tailed student T-Test, Error bars = SEM). **B).** Effect of depletion of PlexinB1, PlexinB2 or RanBPI on nuclear morphology i). Representative images of Hela cell nuclei stained with Dapi, scale bar =10μm. Graph shows average nuclear form factor score for each condition, calculated as 4*π*Area/Perimeter (*p=<0.05, 2 tailed student T-Test, Error bars = SEM). **C).** Representative images of NP1542 cells transfected with control siRNA or two different siRNAs against PlexinB1, stained for tubulin, lamin and Dapi. Scale bar = 10μm. Graph shows % of NP1542 cells with two or more nuclei in cells transfected with control siRNA or siRNA to PlexinB1 (128 or more cells scored per condition per experiment, n=3, *p=<0.01, 2 tailed student T-Test, Error bars = SEM) **D). i)** Representative images of metaphase spreads of 22RV1 cells transfected with control siRNA or siRNA to PlexinB1, Scale bar = 20μm. ii). graph shows % metaphase spreads with indicated number of chromosomes, (average of 3 experiments, 20 or more metaphase spreads counted per experiment per condition. Error bars = SEM), iii). Western blot analysis of knockdown of PlexinB1 by siRNA in 22RV1 cells

In addition, depletion of PlexinB1 in 22RV1 cells (prostate cancer cells with a near normal karyotype) increased the number of metaphase spreads/cells with an abnormal number of chromosomes (Figure 6D).

One indicator of inappropriate exit from mitosis without correct chromosomal segregation is the formation of micronuclei – small nuclear envelope-bound structures in the cytoplasm that contain chromosomes or chromosome fragments. Micronuclei form from lagging chromosomes in anaphase or chromosome fragments, following errors in mitosis or DNA damage ^36^ ^37^. Hela cells or NP1542 cells were treated with Sema4D, a ligand for PlexinB1 and PlexinB2 and the percentage of cells with micronuclei scored. Micronuclei were defined as small DNA-containing structures in the cytoplasm with a nuclear membrane identified by laminA staining. Sema4D treatment resulted a significant increase in the number of Hela and NP1542 cells with micronuclei (Figure 7).

**Figure 7.**
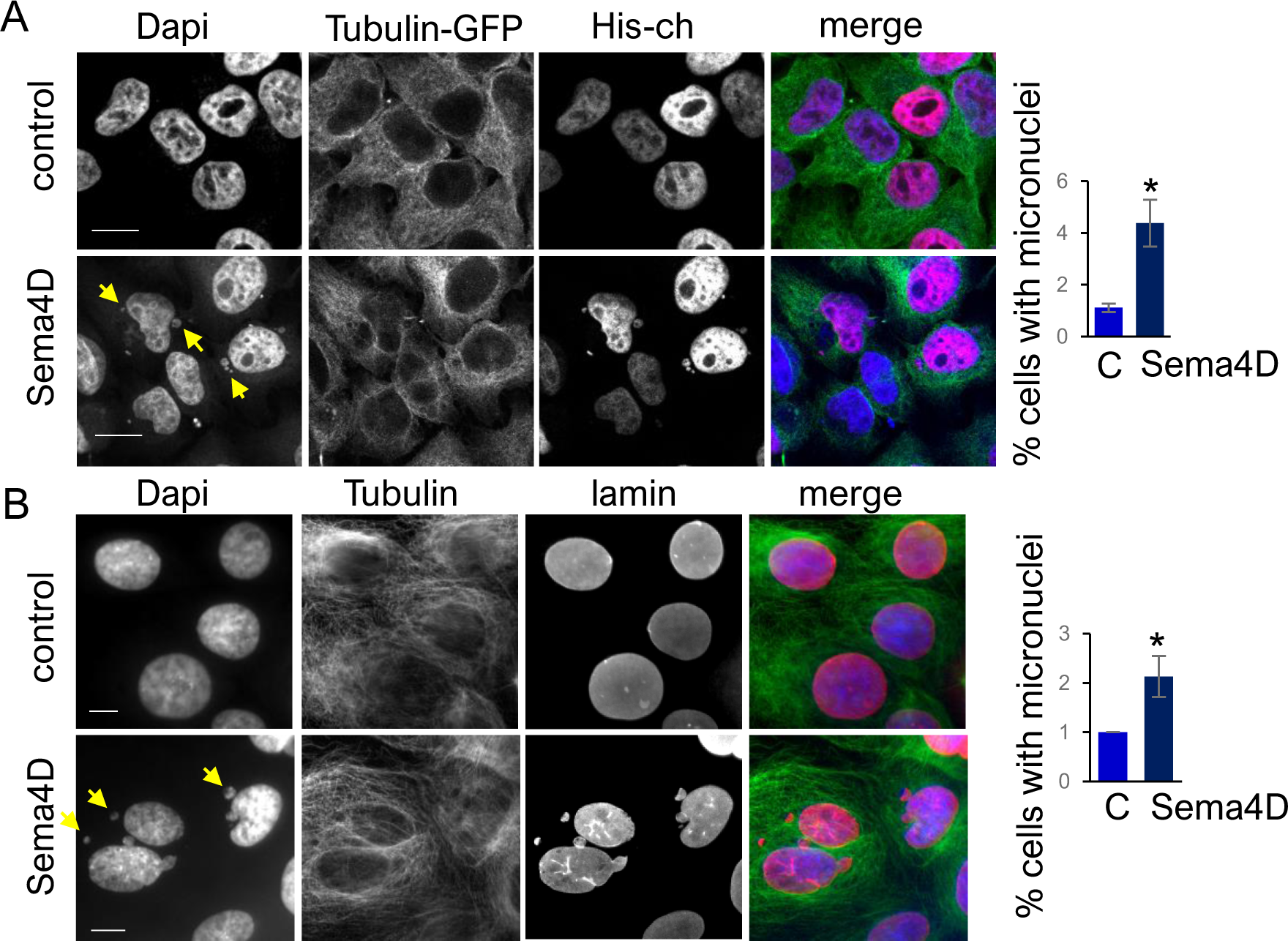
**Activation of PlexinB1/2 with Sema4D increases the % of cells with micronuclei**. **A).** HeLa cells expressing GFP-tubulin and mCherry-histone were treated with Sema4D-Fc (2μg/ml) or vehicle control for 6 hours, then fixed and stained for Dapi (x63 magnification scale bar=10μm) Bar chart shows the % of cells with micronuclei (1750+ cells scored per condition, n=3, *p=<0.05, 2 tailed student T-Test). Error bars = SEM. **B).** NP1542 cells were treated with Sema4D-Fc (2μg/ml) or vehicle control for 6 hours, then fixed and stained for tubulin, lamin and Dapi (x40 magnification scale bar=5μm) Bar chart shows the % of cells with micronuclei (315+ cells scored per condition n=3, *p=<0.05, 2 tailed student T-Test) Error bars = SEM .

Together these results show that PlexinB1 depletion or activation affects chromosomal stability.

## Discussion

Our results have shown that PlexinB1/B2 has a role the regulation of chromosome segregation in mitosis. Mitotic defects resulting from PlexinB1 depletion is restored by inhibition of RCC1, suggesting that Plexins signal via Ran to regulate mitosis.

Mitotic spindle assembly is dependent on the coordinated nucleation of microtubules (MTs), involving centrosome-mediated, chromatin-mediated and MT-mediated MT nucleation pathways in somatic cells. ^23^ ^24^ RanGTP, which is concentrated around the chromosomes due to chromatin-bound RCC1, ^10^ primarily promotes chromosome-dependent spindle microtubule nucleation and stabilisation close to the chromosomes, affecting early stages of spindle assembly. ^13^ ^23^ ^14^ Ectopic expression of constitutively active form of Ran, Ran(Q69L), induces the formation of microtubule asters in xenopus egg extracts^12^ and the cytoplasm of somatic cells ^23^. We found that knockdown of PlexinB1, which increases RanGTP levels^18^, increases acentrosomal microtubule regrowth. At a later time point, depleted cells showed defects in spindle pole focussing. B-type plexins may therefore contribute to spindle assembly by affecting chromosome-dependent spindle assembly.

RanGTP in the vicinity of the chromosomes induces the release of importin-bound spindle assembly factors (SAFs) from importins, facilitating spindle MT nucleation and stabilisation.

Plexins may regulate this process by affecting Ran activity through its function as a RanGAP

Defects in mitosis were seen for PlexinB1 depletion and also upon activation of PlexinB1/2 with Sema4D, which increases or decreases RanGTP levels^18^ respectively; Sema4D treatment led to mitotic cells with collapsed spindles. Cells expressing either constitutively active RanQ69L or the dominant-negative RanT24N give rise to abnormal spindles, showing that the correct balance between RanGTP/GDP is essential for proper spindle formation. ^38^

Defects in spindle formation leads to a delay in metaphase by activation of the SAC ^29^ Depletion of PlexinB1 resulted in an increase in unattached kinetochores, as indicated by Mad1 staining, and a significant increase in the time taken to reach anaphase. Depletion of RanBP1 similarly leads to prolonged metaphase, ^30^ while RanGAP1-depleted MEFs exhibit markedly elongated anaphase and telophase. ^17^

An increase in the percentage of dividing cells with lagging chromosomes and chromosomal bridges was observed in PlexinB1-depleted cells that had bypasses SAC, possibly through merotelic kinetochore attachment.^36^ Similar results have been shown following knockdown of RanGAP1.

In addition to RanGTP expression around chromosomes during mitosis, low levels of RanGTP and RanBP1 are expressed in centrosomes in association with AKAP450 and regulate centrosomal MT nucleation, cohesion, and duplication licensing.^15^

RanGAP1, however, is absent from mt minus ends where mts connect to centrosomes ^31,39^

Our confocal images of digitonin-permeabilised cells suggests that PlexinB1 is expressed around the spindle and also, unlike RanGAP1, at the spindle poles. Depletion of PlexinB1 increased the percentage of cells with multipolar spindles and more than two centrosomes. Expression of GFP-Ran^Q69L^ or GFP-RCC1 mislocalization also increases the number of mitotic cells with multipolar spindles. ^40^ In contrast, an increase in multiple centrosomes has not been reported for RanGAP1 knockdown, although an increase in multipolar spindles is seen upon depletion of RanBP2 which results in the mislocation of RanGAP1^39^. These results suggest that PlexinB1 may serve a function distinct from that of RanGAP1 in centrosomal regulation.

Both the increase in the time to complete mitosis and the increase in multipolar spindle assembly following PlexinB1 depletion was reversed by inhibition of RCC1, implicating Ran in both processes. PlexinB1 may therefore exert its role in mitosis through its activity as a GAP for Ran.

The small RhoGTPases CDC42 and RhoA also have a role in mitosis.^41^ RhoA mediates cell rounding, cortical stiffening and cytokinesis while CDC42 has roles in attachment of spindle microtubules to kinetochores, centrosome integrity and spindle orientation.^41^ B-class plexins regulate several small RhoGTPases, regulating RhoA/C via PDZRhoGEF/LARG^42^ and p190RhoGAP^43^and Plexin-B2 controls mitotic spindle orientation indirectly through CDC42 via its GAP domain.^44^ PlexinB1 may also affect mitosis through these pathways. Plexin-B1 and B3 interact with microtubule end-binding proteins (EBs) and overexpression of the intracellular domains of B-plexins or Sema4D treatment increases the number of MT tip rescues in hippocampal neurons^45^. Thus, PlexinB1/2 could also affect mitosis through EB regulation

Depletion of PlexinB1/2 led to an increase in multinucleate cells and cells with altered nuclear morphology and abnormal karyotype, suggesting that PlexinB1 contributes to chromosome stability.

Furthermore, activation of PlexinB1/B2, by treatment with Sema4D, increased the number of cells with micronuclei, suggesting an inappropriate exit from mitosis. Micronuclei can contribute to genome instability and are a source of chromothripsis^37^ in cancer cells. RanGAP1 depletion promotes micronuclei, chromothripsis and aneuploidy in osteosarcoma ^17^DNA in the cytoplasm can also activate the cGAS-STING pathway leading to expression of proinflammatory genes ^46^

Plexins and their ligands, semaphorins, have a role in cancer and can promote or delay cancer progression depending on cancer subtype.^20,47^ For example, PlexinB1 overexpression is a bad prognostic factor in ErbB2-amplified breast cancer ^48^and ovarian cancer ^49^while Sema3C, a ligand for PlexinB1 is overexpressed in prostate cancer. ^22^ PlexinB1 is mutated in prostate cancer^50^ and Prostate epithelial cell-specific overexpression of a mutant form of PlexinB1 in vivo promotes metastasis ^51^ Conversely loss of Plexinb1 is associated with poor prognosis in melanoma^52^ and estrogen receptor–positive breast cancer. ^51^

Together these results show that defects in B-type plexins may contribute to cancer progression by inducing chromosomal instability.

## METHODS

### Cell culture and transfection

All cell lines were from ATCC (LGC standards, Middlesex, UK) and were STR typed to confirm their identity. HeLa cells were grown in Dulbecco’s modified Eagle’s medium (DMEM) containing 10% foetal calf serum (FCS). NP1542 and 22RV1 cells were grown in RPMI medium with 10% FCS (Invitrogen). Cells were transiently transfected with the indicated plasmids using Lipofectamine 2000 or 3000(Invitrogen) as per manufacturer’s instruction.

### siRNA

Short interfering RNA (siRNA) sequences targeting PlexinB1, PlexinB2 and RanBP1 were obtained from Dharmacon. Plexin expression was knocked down using two different siRNA oligos against PlexinB1: siPlexin B1-1 (*GAGAGGAGCCGACUACGUA*), siPlexin B1-2 (*GCAGAGACCUCACCUUUGA*), against PlexinB2: siPlexin B2-1 (*GCAACAAGCUGCUGUACGC*) siPlexin B2-2 (*UGAACACCCUCGU GGCACU*) or siRNA against RanBP1 (Dharmacon J-006627-06-0005, J-006627-09-0005) together with siGENOME non-targetting siRNA pool (Dharmacon) as control. 25 nM of siRNA was transfected using Dharmafect (Dharmacon) according to manufacturers’ instructions. After 48 h, cells were fixed for immunofluorescence or after 72 h for time lapse video microscopy.

### Expression vectors

pcDNA3-PlexinB1 full-length-myc (PlexinB1-myc), was a kind gift of Dr I Oinuma (Kyoto, Japan)

### Immunocytochemistry

Cells grown on coverslips were fixed (4% paraformaldehyde), permeabilized (0.2% triton), and stained by immunofluorescence. The following primary antibodies were used: mouse anti-Myc antibody clone 9E10 (13-2500 Sigma-Aldrich), rabbit polyclonal anti-Myc (Sigma-Aldrich) Mouse anti-tubulin (clone DMIA, Sigma-Aldrich), rabbit anti-pericentrin (Abcam), anti-lamin A antibody produced in rabbit (Sigma-Aldrich L1293-200UL), Mad1 (GeneTex, GTX105079), Alexa-Fluor488, Alexa-Fluor555, Alexa-Fluor594, or Alexa-Fluor647- conjugated secondary antibodies and Alexa-Fluor 633 Phalloidin were from Life Technologies. Coverslips were mounted with Prolong Gold Antifade mountant (Life Technologies). Images were taken at x63 magnification using a Zeiss LSM510 confocal microscope or at 40x or 60x magnification using a Nikon A1 inverted confocal with spectral detector or a Nikon Inverted spinning disk confocal microscope, or an Olympus IX71 inverted microscope.

### Sema4D treatment for micronuclei studies

HeLa cells expressing GFP-tubulin and mCherry-Histone or NP1542 cells, plated on coverslips, were treated with Sema4D-Fc(2g/ml, Bio-techne) for 6 hours, then fixed and stained for Dapi and pericentrin (Hela cells) or laminA, Dapi and pericentrin (NP1542 cells). Images were taken at x40 magnification using an Olympus IX71 inverted microscope. Micronuclei were scored by counting the number of cells with extracellular DNA identified by Dapi, separate from the nucleus, with a concomitant gap in tubulin staining, or encapsulated with lamin ImageJ was used to score the percentage of cells with micronuclei.

### Digitonin treatment

Cells plated on coverslips were washed with PBS and then treated with 0.005% digitonin (Sigma) in 110mMKOAc, 20mM Hepes pH7.3, 2mMMgOAc, 0.5mM EGTA, 2mM DTT, 1ug/ml aprotinin, 1ug/ml leupeptin (Sigma) for 4mins, washed with PBS and fixed with 4%PFA.

### Apoptosis assay

HeLa cells expressing GFP-tubulin plated on coverslips were transfected with non-silencing siRNA or two different siRNAs against PlexinB1. 48hrs after transfection the cells were fixed and stained for activated Caspase-3/7 using Apoptosis CellEvent™ Caspase-3/7 Green Detection Reagent (Thermo Fisher Scientific) according to manufacturers’ instructions. The percentage of cells stained for activated caspase 3/7 were scored using imageJ (n=3 a total of 3373 or more cells were scored per condition).

### Time lapse microscopy

HeLa cells expressing tubulin-GFP and Histone-cherry plated in chamber slides were transfected with siRNA to PlexinB1 or control non-silencing siRNA. 72 h after siRNA transfection the medium was changed to FluoroBrite DMEM Media (Thermo Fisher Scientific) with 10% serum, 1mM sodium pyruvate and 2mM glutamine. The cells were imaged using a Nikon Inverted spinning disk confocal microscope every 5 mins and videos created using NIS-Elements imaging software. The length of time of each dividing cell to go from prometaphase to anaphase, the percentage of cells with lagging chromosomes in anaphase and the percentage of cells with multipolar spindles were scored from the videos. For rescue experiments with the RCC1 inhibitor, cells transfected with siRNA to PlexinB1 or control non-silencing siRNA were treated with EPZ020411 (50m) or DMSO control 5 hours before the 18hr timelapse videos were started. 100+ cells were scored per condition per experiment and the experiments were repeated independently 3 times.

For videos of Sema4D-Fc-647 endocytosis, HeLa cells expressing GFP-tubulin and mCherry-Histone plated in chamber slides were incubated with Sema4D-Fc-Alexa647 and imaged using a Nikon Inverted spinning disk confocal microscope and imaged at x40.

### Immunoblotting

For immunoblotting, proteins were resolved by SDS-PAGE and transferred to nitrocellulose membrane (Schleicher and Schuell). Membranes were blocked with TBS (20 mM Tris-HCl [pH 7.6], 137 mM NaCl) containing 5% non-fat dried milk and 0.05% Tween 20. Bound antibodies were visualised with horseradish peroxidase-conjugated goat anti-immunoglobulin G antibodies and enhanced chemiluminescence (ECL; Amersham Pharmacia Biotech). The following primary antibodies were used: anti-PlexinB1 (ECM Biosciences), anti-PlexinB2 (R&D Systems), Mouse anti-GAPDH (MAB374, Millipore, now Merck), Secondary antibodies used for Western blotting: horse-radish peroxidase linked (HRP) polyclonal goat anti-mouse IgG (P0447), goat anti-rabbit IgG (P0448), rabbit anti-goat IgG (P0449) (Agilent) or goat anti-rat IgG (sc-2006) (Santa Cruz Biotechnology).

### Microtubule regrowth assay

Hela cells expressing tubulin-GFP and Histone-cherry, grown on coverslips were transfected with non-silencing siRNA or two different siRNAs against PlexinB1. 48hrs later, the cells were incubated on ice for 30mins. The medium was replaced with prewarmed (37°C) DMEM+10% FCS and the cells incubated for 0-30min at 37°C. Cells were fixed at various time points and stained for pericentrin.

### Cell synchronisation

HeLa cells expressing expressing tubulin-GFP and Histone-cherry, transfected with siRNA to PlexinB1, PlexinB2, RanBP1 or control non-targeting siRNA were treated with 10 µM RO-3306 in DMSO or DMSO as control for 20 hours.

Cells were then washed twice with the fresh media, then cultured further with the fresh media and fixed at various time intervals.

Fixed cells were then stained with DAPI and mounted on slides for imaging.

Cells at prometaphase, metaphase, anaphase, telophase and cytokinesis were counted at each time interval.

### Karyotype

22RV1 cells were treated with 100ng/ml Colcemid at 37 °C for 45 min. Cells were then washed in PBS, and treated with TrypLE Express (gibco 12604-013) to detach them. The cell suspension was centrifuged at 200 x g for 10 min. and the pellet resuspend in 0.075 M KCl prewarmed to 37°C, vortexed, then incubated at 37°C for 10 min.

Cells were recentrifuged at 200 x g for 5 min at 25 °C and the pellet resuspended in fresh Carnoy’s Fixative (3:1 ratio of methanol:glacial acetic acid) while vortexing. The Carnoy’s Fixative was replaced twice by centrifugation at 200 x g for 5 min. The cells were then dropped onto slides at an angle together with more fixative and stained with Dapi when dry.

### Nuclear morphology

HeLa cells expressing GFP-tubulin and mCherry-Histone plated on coverslips were transfected with non-silencing siRNA or two different siRNAs against PlexinB1, PlexinB2 or RanBP1. The cells were fixed, stained for DAPI and the Nuclear Form factor (calculated as 4*π*Area/Perimeter) of nuclei scored using Cellprofiler (https://cellprofiler.org).

### Statistics

Statistical analysis was performed using Microsoft Excel or GraphPad. Error bars on graphs denote the average values +/- standard error of the mean (SEM), statistical significance was determined by two tailed paired students’ t-test unless otherwise stated.

## Supporting information

Supplimentary images

---

**Supplementary Figure 1.**
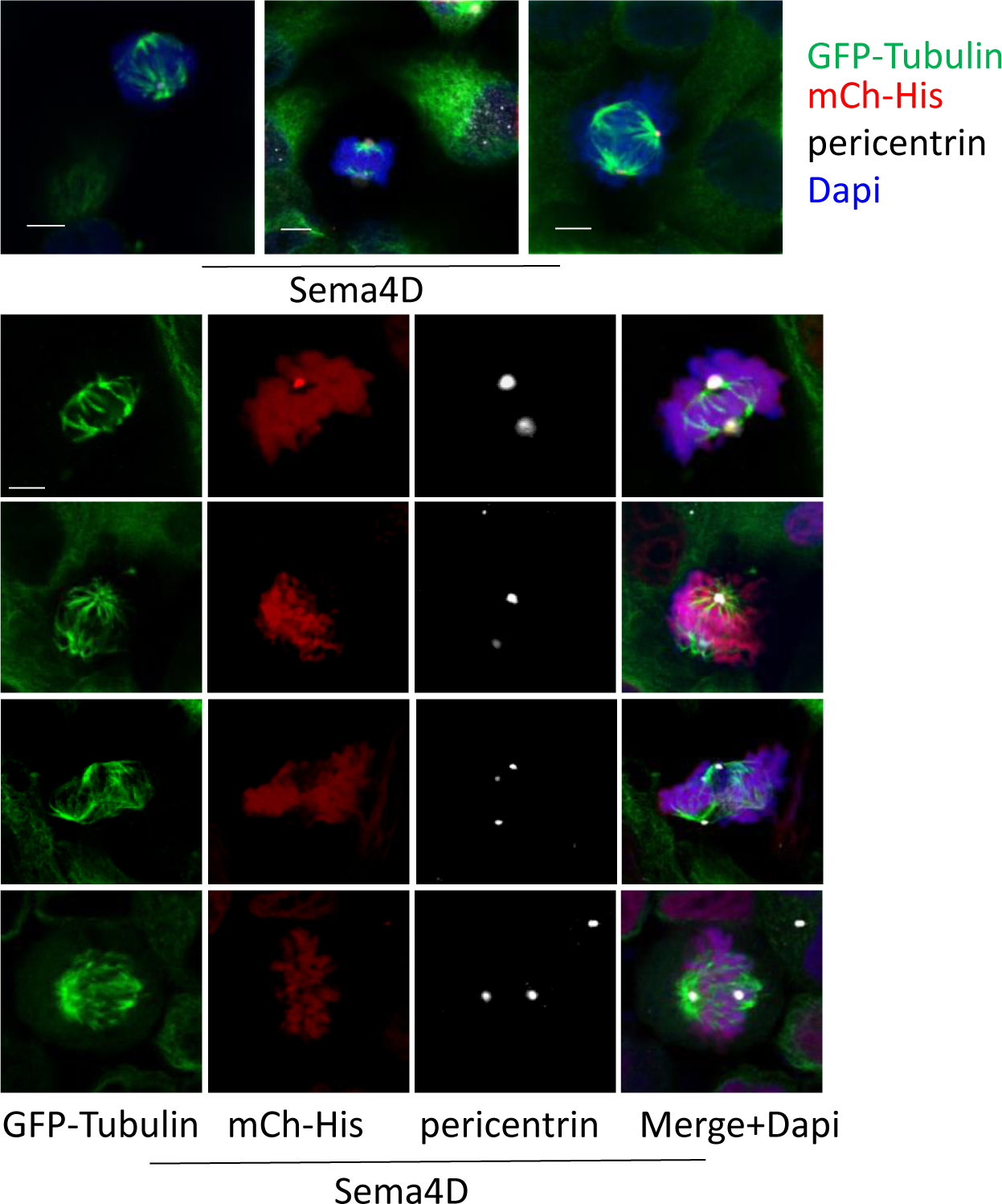
Representative images of dividing HeLa cells expressing GFP-tubulin and mCherry-histone, treated with Sema4D-Fc (2μg/ml) for 6 h, stained for pericentrin and Dapi (x63 magnification, scale bar=5 μm).

**Supplementary Figure 2.**
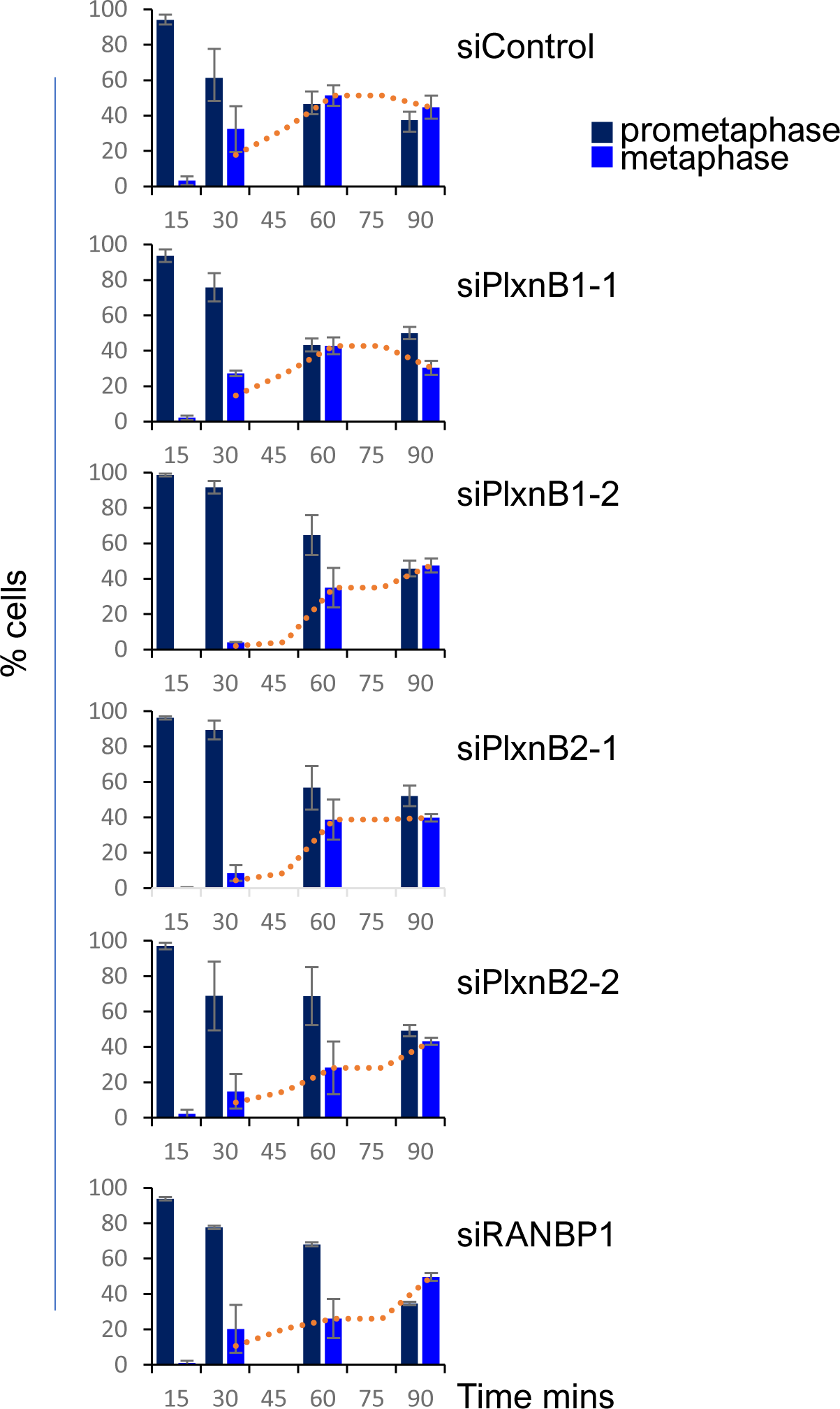
Plexin depletion delays mitosis in G2/M synchronised HeLa cells. Hela cells were synchronised by treatment with 10 μM RO- 3306. The percentage of fixed cells in prophase and metaphase were scored from confocal images at time points indicated, following release from the G2/M block

**Supplementary Figure 3.**
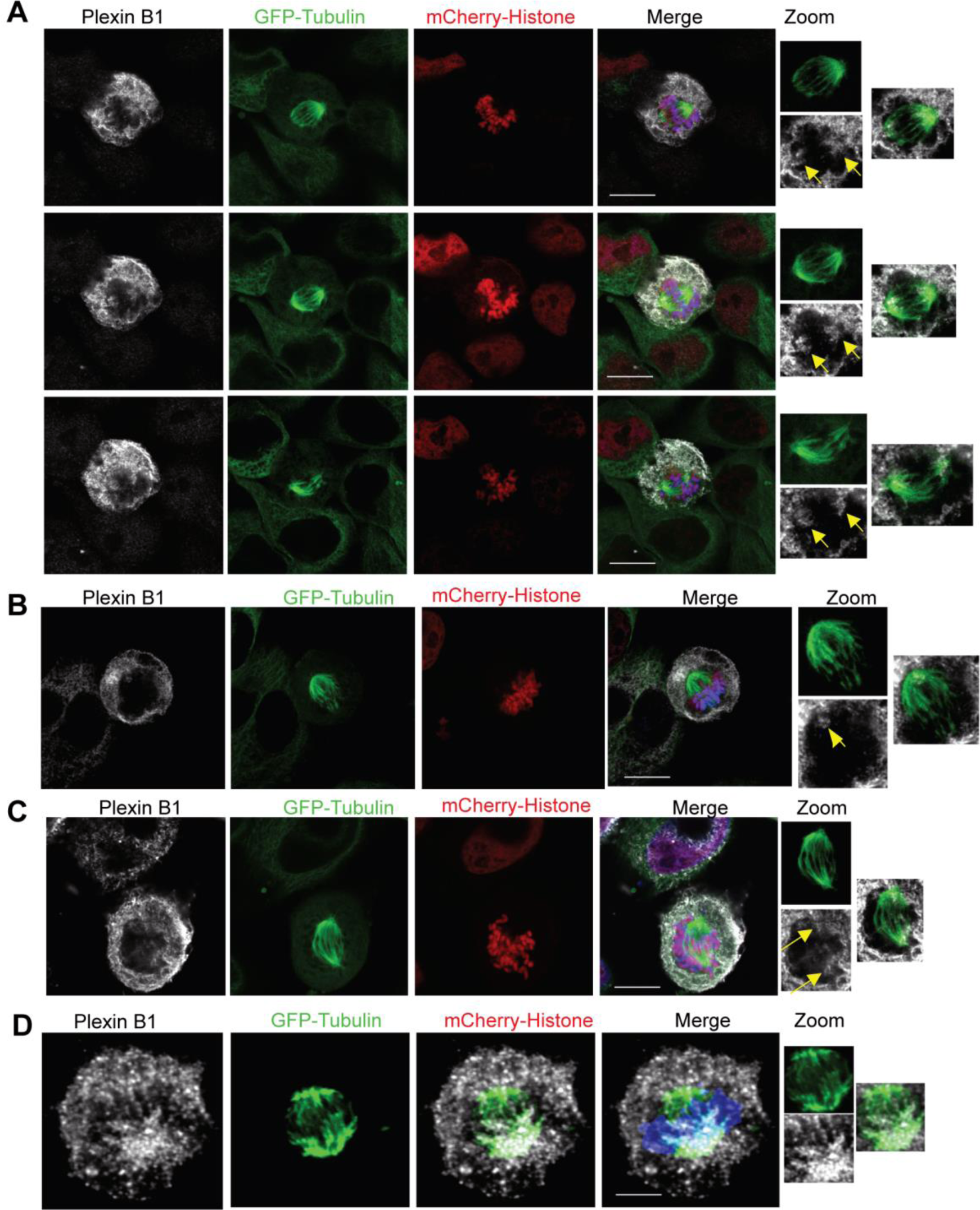

## Supplementary videos

**Supplementary video 1.** HeLa cells expressing GFP-tubulin and mCherry-histone treated with Sema4D-Fc-Alexa647 (white), 2 min time frames, total time of 25 min, x60 magnification

**Supplementary video 2.** HeLa cells expressing GFP-tubulin and mCherry-histone transfected with control siRNA, (control for video 3). 5 min time frames, total time of 150 min, x40 magnification

**Supplementary video 3.** HeLa cells expressing GFP-tubulin and mCherry-histone transfected with siRNA to PlexinB1, (time lapse video taken at same time as Supplementary video 2). 5 min frames, total time of 150 min, x40 magnification,

**Supplementary video 4.** HeLa cells expressing GFP-tubulin and mCherry-histone transfected with control siRNA, (control for video 5). 10 min frames, total time of 90 min, x60 magnification

**Supplementary video 5.** HeLa cells expressing GFP-tubulin and mCherry-histone transfected with siRNA to PlexinB1, 10 min time frames, total time of 200 min, x60 magnification.

