## Supplementary material for "B-type plexins regulate mitosis via RanGTPase": Supplimentary images

### Supplementary Figure 1

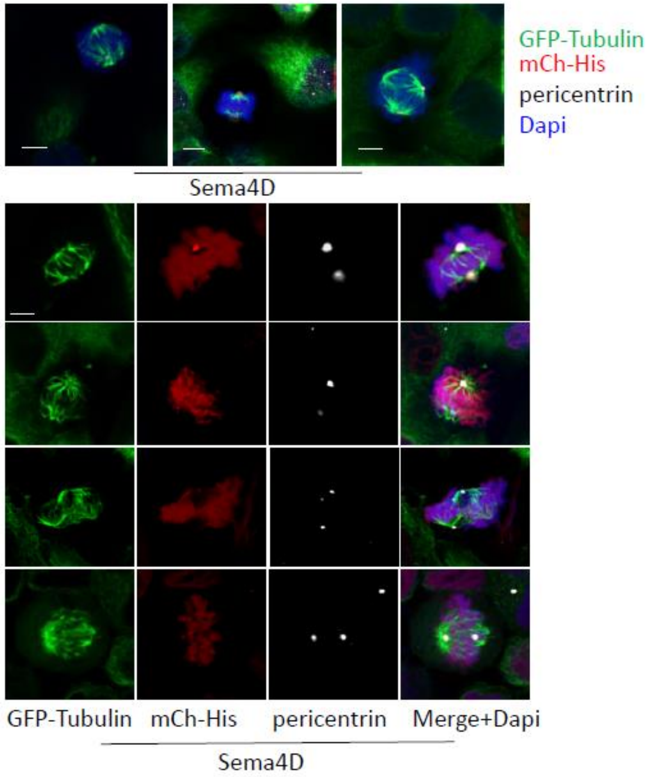


Supplementary Figure 1

Representative images of dividing HeLa cells expressing GFP-tubulin and mCherry-histone, treated with Sema4D-Fc (2µg/ml) for 6 h, stained for pericentrin and Dapi (x63 magnification, scale bar=5 µm).

SUPPLEMENTARY FIGURE 2

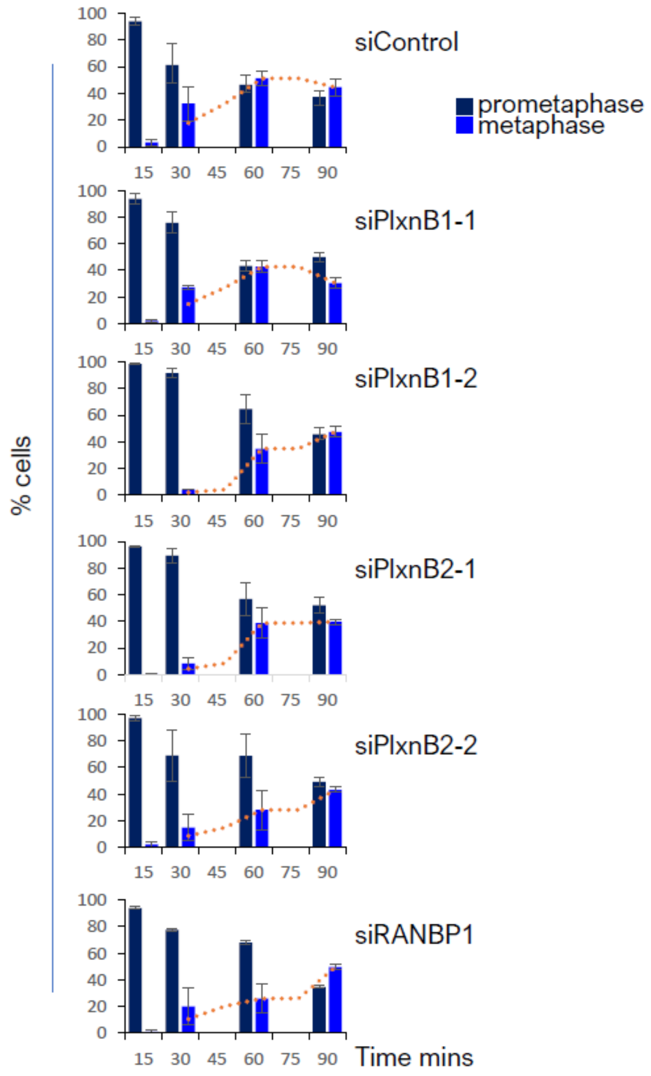


**Supplementary Figure 2 Plexin depletion delays mitosis in G2/M synchronised HeLa cells**
Hela cells were synchronised by treatment with 10 μM RO-3306. The percentage of fixed cells in prophase and metaphase were scored from confocal images at time points indicated, following release from the G2/M block

Supplementary 3

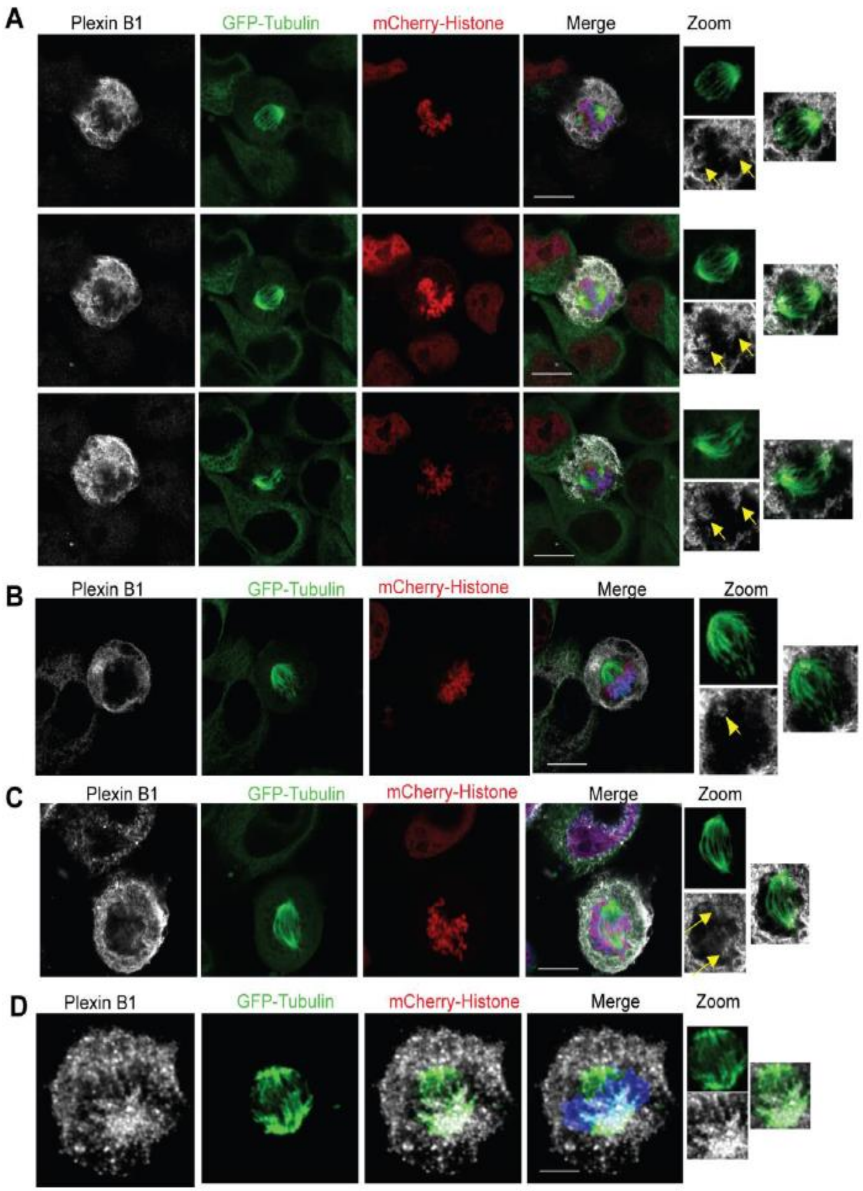
